# Recognizing the Continuous Nature of Expression Heterogeneity and Clinical Outcomes in Clear Cell Renal Cell Carcinoma

**DOI:** 10.1101/044891

**Authors:** Xiaona Wei, Yukti Choudhury, Weng Khong Lim, John Anema, Richard J. Kahnoski, Brian Lane, John Ludlow, Masayuki Takahashi, Hiro-omi Kanayama, Arie Belldegrun, Hyung L. Kim, Craig Rogers, David Nicol, Bin Tean Teh, Min-Han Tan

## Abstract

**PURPOSE:** Evaluation of 12 ccRCC publicly-available ccRCC gene expression datasets showed that previously proposed discrete molecular subtypes are unstable. To reflect the continuous nature of gene expression observed, we developed a quantitative score (Continuous Linear Enhanced Assessment of Renal cell carcinoma, or CLEAR) using expression analysis founded on pathologic parameters.

**MATERIALS AND METHODS:** 265 ccRCC gene expression profiles were used to develop the CLEAR score, representing a genetic correlate of the continuum of morphological tumor grade. A signature derivation method based on correlation of CLEAR score with gene expression ranking was used to derive an 18-transcript signature. External validation was conducted in multiple public expression datasets.

**Results:** As a measure of intertumoral gene expression heterogeneity, the CLEAR score demonstrated both superior prognostic estimates (94% vs 83% adequacy index in TCGA dataset) and inverse correlation with anti-angiogenic tyrosine-kinase inhibition (65% vs 55% adequacy index) in comparison to previously proposed discrete subtyping classifications. Inverse correlation with high-dose interleukin-2 outcomes was also observed for the CLEAR score (p=0.05). Multiple somatic mutations (VHL, PBRM1, SETD2, KDM5C, TP53, BAP1, PTEN, MTOR) were associated with the CLEAR score. Application of the CLEAR score to independent expression profiling of intratumoral ccRCC regions demonstrated its ability to reflect intratumoral expression heterogeneity and further analysis showed average intertumoral heterogeneity exceeded intratumoral heterogeneity.

**Conclusions:** The CLEAR score, a gene expression signature developed on histopathology, outperformed discrete subtype-classification in prognostic estimates and correlated better with treatment outcomes. Recognizing cancer as a continuum has important implications for laboratory and clinical research.

## Introduction

Clear cell renal cell carcinoma (ccRCC) is a heterogeneous disease with diverse morphologies, molecular characteristics, clinical outcomes and therapeutic responses [1,2]. Clinicopathologic prognostic factors including tumor stage, nuclear grade, morphologic characteristics, tumor size, nodal status and patient performance status [3-5] have been studied as features to classify and predict disease-specific survival. However, these prognostic factors have limited accuracy for tumors of intermediate grade and stage that form a substantial proportion of ccRCC cases.

To improve prognostic prediction, several studies have proposed dividing ccRCC into molecular subtypes, primarily through unsupervised clustering of gene expression profiles or genetic alterations. Although these methods were reported to yield good performance in predicting survival, the results may be susceptible to instability arising from the analytic techniques used [6,7]. There is also insufficient rigorous testing and evaluation of the performance of these subtype classification approaches in independent datasets [8-15].

In this study, we examined 12 ccRCC datasets using consensus clustering sensitivity analysis by varying key parameters including item, feature, distance and iteration. We found that the clustering techniques yielded relatively unstable sample classification. Based on the observed lack of clear delineation between prognostic subtypes, we recognized that a genetic continuum would possibly reflect underlying biologic, histopathologic and clinical heterogeneity better than subtyping. To reflect this probable underlying reality, we developed an alternative strategy involving a continuous quantitative expression assessment of ccRCC tumor prognosis using tumor grade, which is a key histopathologic parameter in determining tumor behavior across different cancers. We thus created a continuous CLEAR score (continuous linear enhanced assessment of ccRCC) based on 18-transcript signature derived from a large internal dataset. In applying it to multiple external datasets, we investigated the performance of the CLEAR score, as a continuous measure of intertumoral expression heterogeneity, in estimating survival outcomes as compared to subtyping approaches. We evaluated outcomes of anti-angiogenic tyrosine-kinase inhibition, as well as high-dose interleukin-2 (IL-2) therapy in relation to this continuous CLEAR score. We further applied the CLEAR score to investigate 65 heterogeneous tumor regions from 10 ccRCC patients to determine extent and importance of intratumoral expression heterogeneity. The CLEAR score package is available at: https://sourceforge.net/user/verification?hash=3dfcf804138b6b959a315335e0203a19

## Results

### Evaluation of ccRCC subtyping with public datasets

To investigate the performance of ccRCC subtyping, 12 public datasets were obtained from the GEO database, EMBL-EBI and The Cancer Genome Atlas (TCGA) (Supplementary Table S1). Sensitivity analysis of consensus clustering done through varying key parameters demonstrated a discrepant number of predicted clusters ranging from 2 to 4 (Supplementary Table S2). Analysis by principal component analysis (PCA) showed poor delineation of proposed molecular subtypes (Supplementary Figure S1).

### Assessment of 265 internal ccRCC samples with the CLEAR score

In light of the above results, we hypothesized that a continuous scale might be more representative of tumor biology. The CLEAR score algorithm (Supplementary Figure S2) was thus designed. With our internal 265 ccRCC gene expression profiles, the continuous scale were derived by calculating the correlation distance between the sample of interest and two reference sample sets (RSSs) with distinct tumor grade (RSS1 with tumor grade 1 sample sets and RSS2 with tumor grade 4 sample sets).

This scale was then normalized (range from 1 to 100) and defined as CLEAR score (Supplementary Dataset S1). We determined that various clinical variables correlating with biological aggressiveness such as grade, stage and size distributed continuously across the CLEAR score (Figure 1A). Sarcomatoid histology of ccRCC are correlated with higher CLEAR score, with 9 of 14 samples having a score exceeding 80 (Supplementary Table S3). Corresponding survival analysis of these 265 ccRCC yielded varying cancer-specific survival outcomes across the CLEAR score (p<1e-02) (Figure 1B).

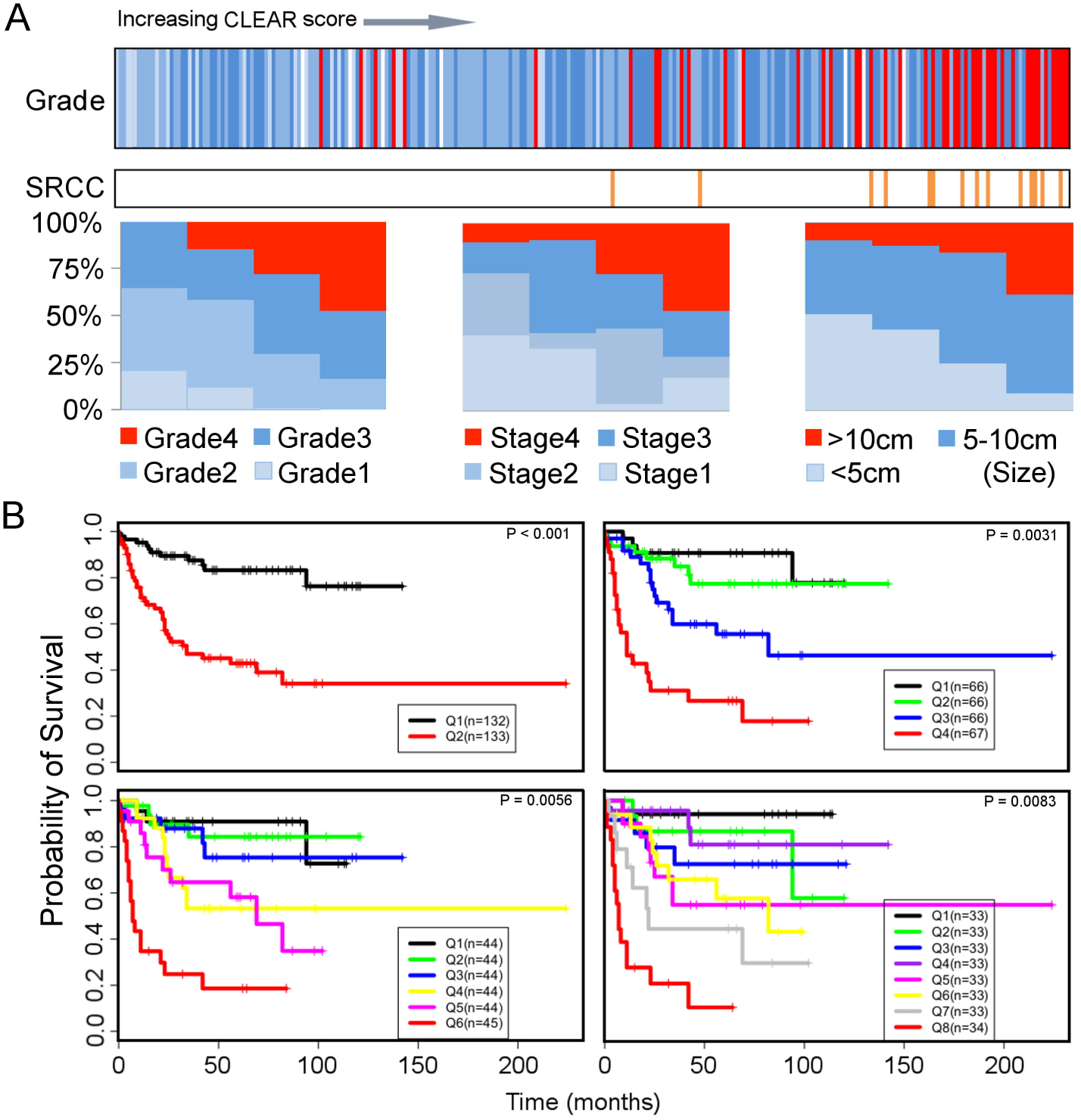

### Derivation of molecular drivers of CLEAR score and validation in TCGA datasets

A signature derivation method based on correlation of CLEAR score with gene expression was developed (Supplementary Figure S3), yielding an 18-transcript signature. To independently evaluate the performance of the CLEAR score, we applied it to the TCGA-414 cohort [15] generated by the RNA-seq. The expression profiles of 18-transcript signature exhibited a similar expression pattern between our internal dataset and independent TCGA dataset as presented in Figure 2. An association between tumor advanced grade, stage and size with high CLEAR score were observed in Figure 3A with TCGA dataset. Kaplan-Meier survival analysis presented a significant difference in cancer-specific survival among gradient groups (p<1e-02) (Supplementary Figure S4). We further evaluated 11 additional public ccRCC datasets using the same approach and CLEAR score for each sample are available upon request.

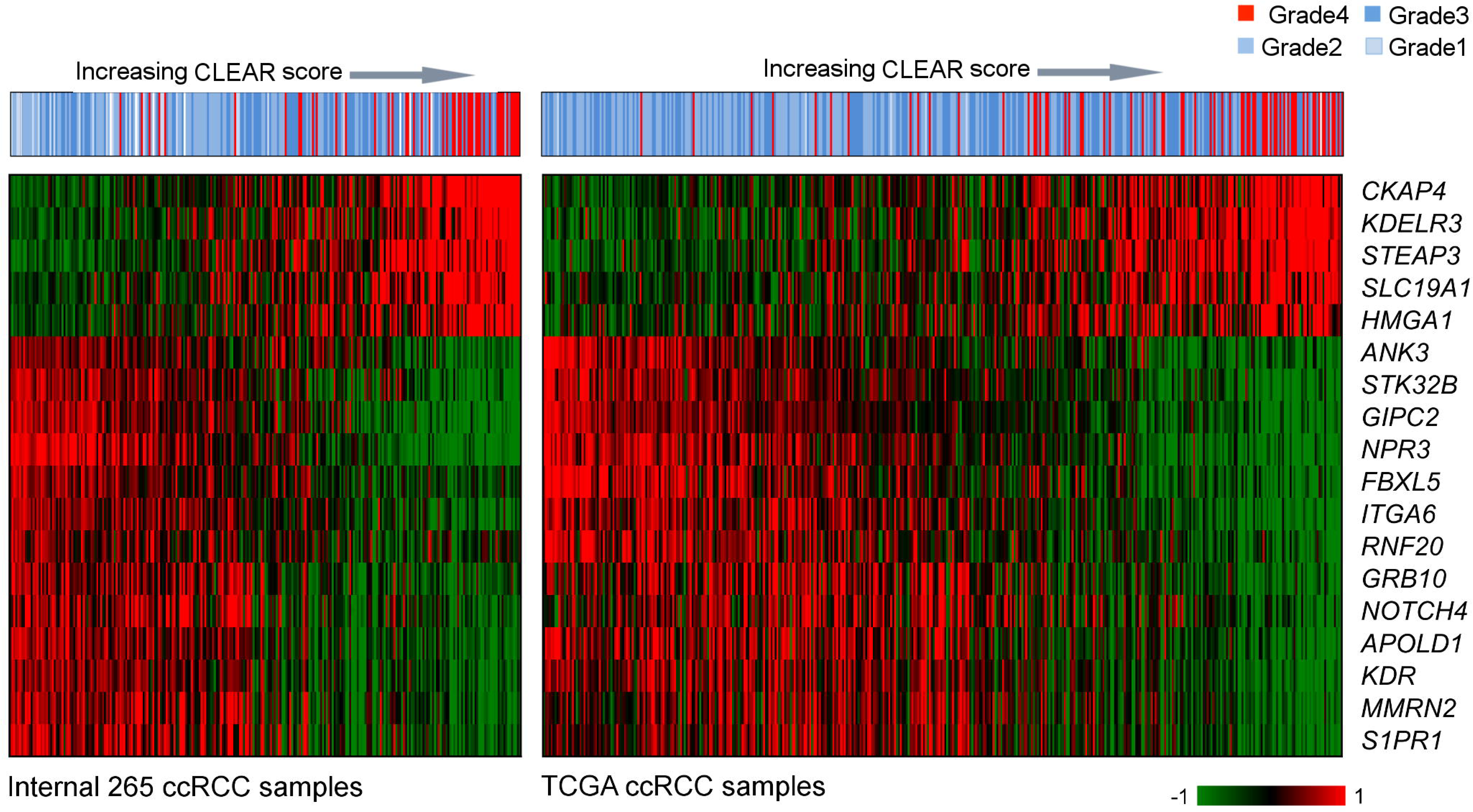

### Comparison of CLEAR with the prognostic subtype model

Two sets of gene expression signatures (110 transcripts and 34 transcripts respectively) have been previously identified for prognostic classification [13,16,17]. To evaluate the prognostic accuracy of CLEAR score for survival outcomes, we compared the predictive value of CLEAR score model with these two prognostic models of CCA/ CCB classification^16,17^ using the independent TCGA dataset. Likelihood ratio testing [18] demonstrated a significant benefit in prediction when adding the CLEAR score model to these two CCA/B subtyping models (110 transcripts and 34 transcripts respectively) in cancer-specific survival tested, with no significant benefit using the converse approach (CSS: p=0.00036 vs. p= 0.059; CSS: p=0.00027 vs. p=0.023). The higher adequacy index of the CLEAR model demonstrated that the expression-based continuous CLEAR score provide improved performance (Figure 4A, Figure 4B).

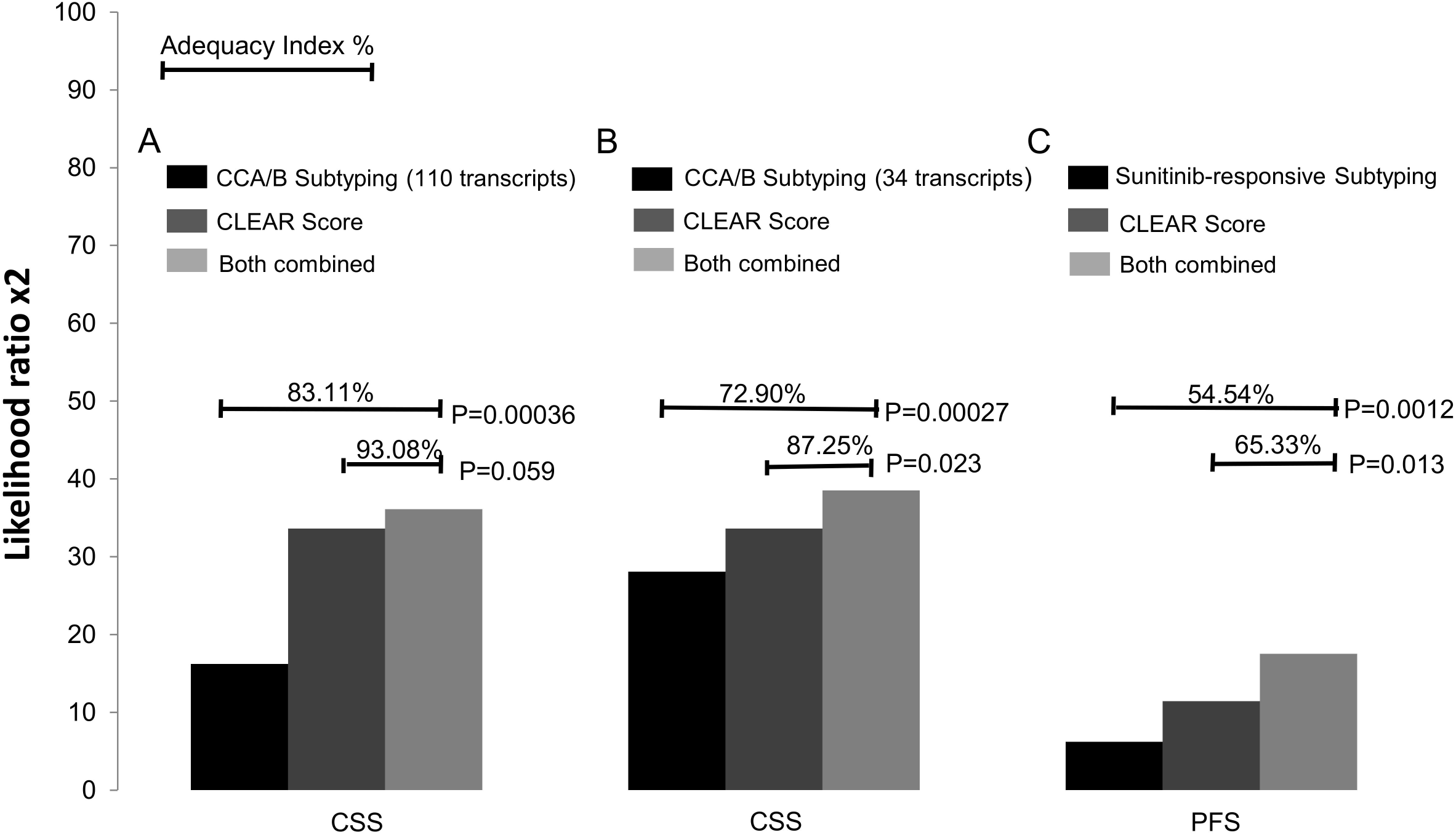

### Association of CLEAR score with therapeutic response

To investigate the potential utility of the CLEAR approach in predicting patient benefit from TKI treatment, we applied CLEAR to a public dataset (E-MTAB-3267) with 53 metastatic French ccRCC patients who received first-line sunitinib[19,20]. Patients who experienced a complete or partial response (CR/PR) presented a relatively lower CLEAR score than patients with progressive disease (PD), who presented with a very high median score (Mann–Whitney U test, p=0.000149) (Supplementary Figure S5). We further compared the predictive value of CLEAR with the model reporting association of subtypes with sunitinib response. Likelihood ratio testing demonstrated a superior performance of the CLEAR score to the proposed molecular subtyping in predicting outcomes (PFS: p=0.0012 vs p=0.013) (Figure 4C).

It is notable that several patients in the internal dataset received high-dose IL-2 treatment, of which 4 and 6 patients experienced durable complete response and eventual progressive disease respectively. High dose IL-2 is associated with durable responses in metastatic ccRCC[21] but is associated with significant toxicities. In general, all patients undergoing high-dose IL-2 therapy had relatively high CLEAR scores (above 50). While sample numbers were low, there was a trend to lower CLEAR scores in patients experiencing complete responses (Mann–Whitney U test, p=0.05) (Supplementary Table S4).

### Assessment of Intratumoral heterogeneity

The CLEAR score can be applied on different regions of the same tumor. We investigated intratumoral heterogeneity using the GSE53000 dataset from GEO database, which contained 65 regions from 10 ccRCC patients [22,23]. Supplementary Dataset S2 presents the CLEAR score for all 65 regions. The boxplot shows the distribution of CLEAR score which indicated that when the median CLEAR score increase, the subsamples tend to have CLEAR score that are more divergent (one sample exception of EV003). Median absolute deviation (MAD) of CLEAR score in each patient was used as an indicator for measurement of the variation of heterogeneity (Supplementary Table S5). To examine intertumoral and intratumoral prognosis heterogeneity, gene expression profiles of the 18-transcript signature from 65 regions of 10 individual tumors[22,23] and 414-TCGA samples were summarized into a heatmap and all the samples ordered based on CLEAR score (Figure 5). The MAD of CLEAR score of all these 65 regions is much larger than the regions within individual tumors, suggesting that intertumoral expression heterogeneity commonly exceeds intratumoral expression heterogeneity. EV005, RMH008, EV006, EV003 have the lower degree of intratumoral heterogeneity while EV001, EV007, RK26, RMH002, EV003, RMH004 has the higher degree of intratumoral heterogeneity (Figure 5, Supplementary Table S5).

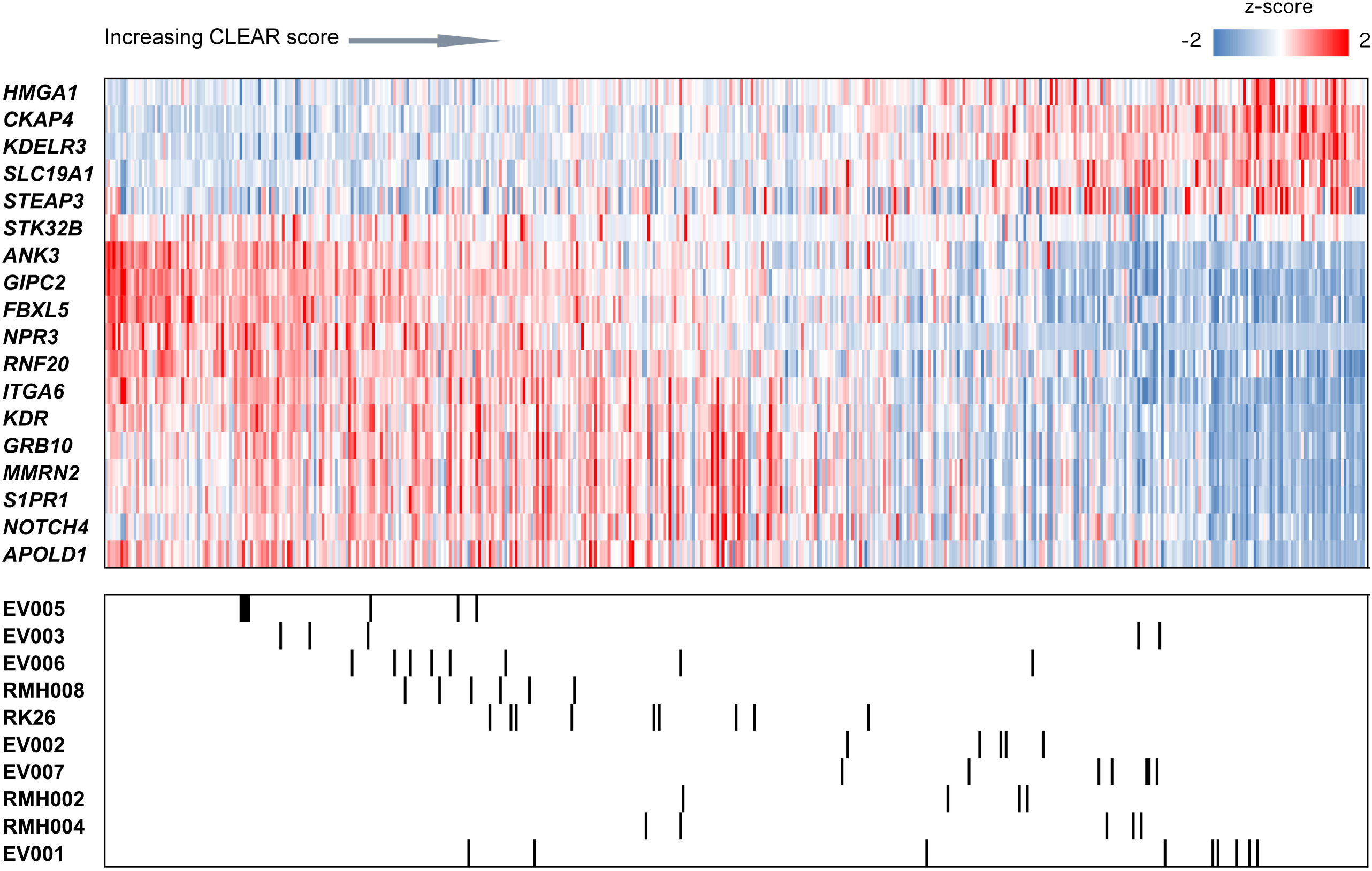

### Correlation of CLEAR score with somatic mutations

To investigate the correlation of CLEAR score with intertumoral mutational heterogeneity, after identifying the CLEAR score of TCGA cohort, we examined the reported ccRCC somatic mutation events per sample (Figure 3A) and found that *BAP1, SETD2, KDM5C,TP53, PTEN* and *MTOR* were associated with higher CLEAR scores (Mann–Whitney U test, *BAP1* (p = 2.32e -08); *SETD2* (p= 0.07); *KDM5C* (p=0.08); *TP53* (p= 0.0004); *PTEN* (p=0.009); *MTOR* (p=0.04)). In contrast, an association between lower CLEAR scores and somatic mutations of *VHL (p=0.09) and PBRM1* (p=0.045) were noted (Figure 3B).

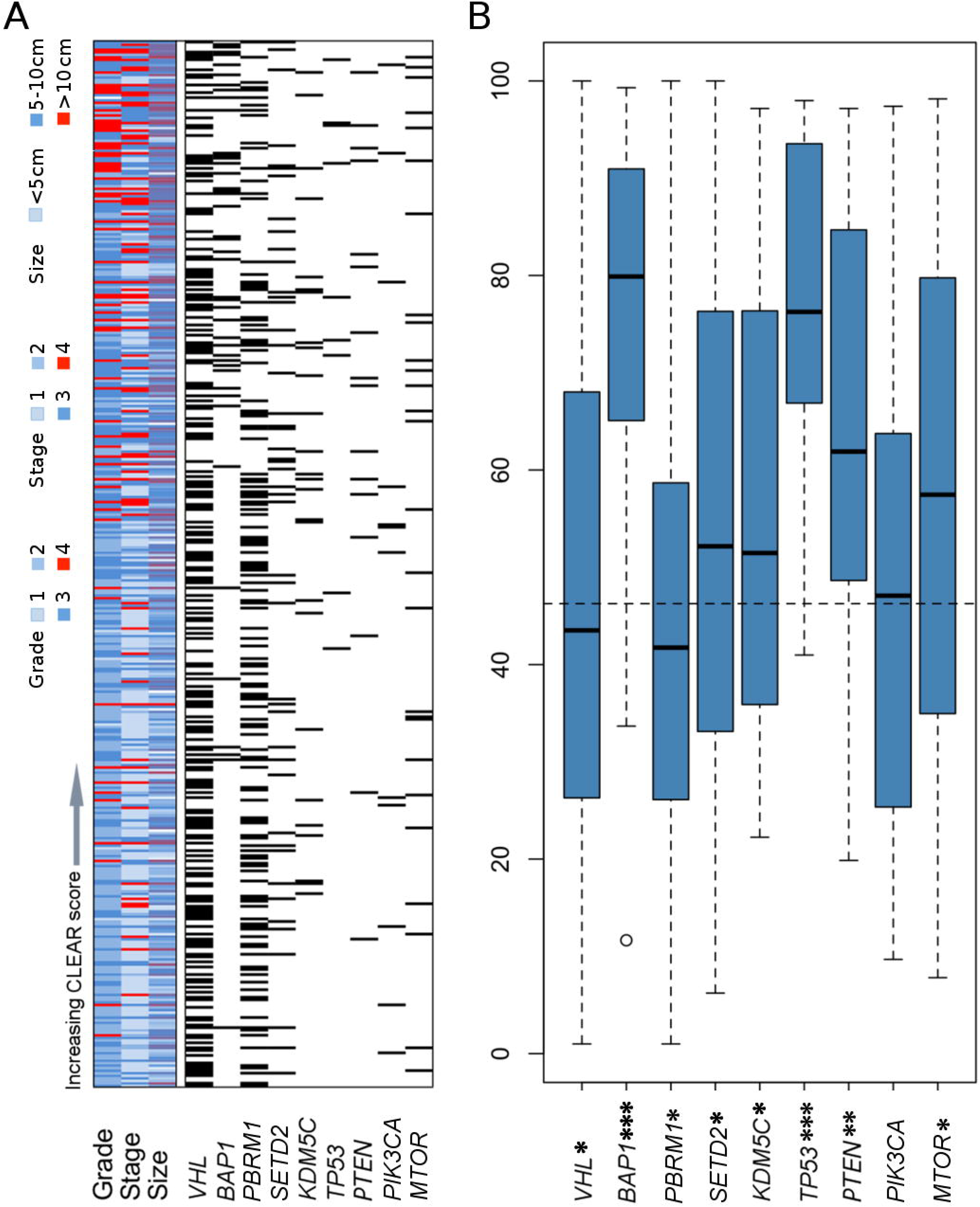

We further evaluated the correlation of CLEAR score with mutation in intratumoral mutational heterogeneity using the GSE53000 dataset [22,23]. A sample region tree with the driver mutation information was constructed using the maximum parsimony method (Supplementary Figure S7). The phylogenetic tree was then annotated with driver gene mutation status and CLEAR score. From the region tree, we can observe most sample regions from the same tumor clustered together with several exceptions caused by intratumoral heterogeneity, which indicates that the degree of intertumoral heterogeneity is larger than that of intratumoral heterogeneity. Our result also showed that *PBRM1* is associated with regions having low CLEAR score, while *SETD2, BAP1* and *KDM5C* are associated with the sample regions which have higher CLEAR score.

## Discussion

In our evaluation of 12 independent datasets, molecular subtyping as derived by unsupervised methods yielded varying subtypes depending on parameters. To reflect the apparent continuum as observed in data analysis, the CLEAR score was designed as a continuous quantitative score derived from histopathologic tumor grade, and corresponding to tumor aggressiveness of ccRCC. Its application to samples allows for evaluation of intertumoral heterogeneity on a continuous scale. Investigation of its performance in multiple independent datasets shows better predictions of prognosis and drug response in comparison to proposed subtyping, suggesting that intertumoral heterogeneity as measured by CLEAR scoring on a continuum is biologically and clinically meaningful. It is of considerable interest that the CLEAR score here, founded on a relatively simple pathology-based morphologic approach, outperformed complex supervised and unsupervised analyses, implying that careful consideration of appropriate pathologic data for integrated analyses may be of meaingful benefit and insights. We speculate that a broader application of this approach to other cancer types may yield helpful insights. The CLEAR score may either be interpreted as a varying mixture of a composite of cells on two extremes of biological aggressiveness, or a clonal expansion of cells that vary across expression heterogeneity. While evaluation of matching CLEAR scores in different regions of the same tumor may help to address such questions, single-cell expression analysis would allow a definitive conclusion.

Gene expression and deep sequencing analysis have provided insights into clinical heterogeneity [22,23]. Corresponding reports have suggested heterogeneous expression patterns with good and poor prognosis signatures coexisting in 8 of 10 cases [16]. Research into intratumoral heterogeneity and its corresponding real-world clinical implications such as drug resistance, are expanding rapidly. The use of the CLEAR score can provide a strong basis for quantifying such variation, thus supporting development of improved diagnostics and therapeutics. Our work has provided direct empirical support for an observation that average intertumoral bulk expression heterogeneity exceeds intra-tumoral heterogeneity, which has not been previously demonstrated. Additional research investigating the origin of such apparent heterogeneity in clonal dynamics or deriving from an ancestral ccRCC clone will have profound implications for drug development. Indeed, we speculate the observed association between improved drug response and lower CLEAR score, may reflect improved outcomes in tumors with lower heterogeneity.

The distinction between a prognostic and predictive biomarker of treatment is important. Beyond a correlation with prognostic outcomes in patients with localized disease, the CLEAR score exhibits an inverse correlation with outcomes of high-dose IL-2 treatment and anti-angiogenic tyrosine kinase inhibition (TKI) in patients with metastatic RCC.

This indicates that the CLEAR score is likely to be a predictive biomarker, the nature of which is best tested in a prospective clinical trial. Given the anti-angiogenic and immune-related mechanisms of each drug class, research to dissect the immune and stromal contributions to tumor gene expression may shed more light on this association.

The limitations of this study are its retrospective design that might influence sample selection, and the relatively limited number of subjects for IL-2 and TKI response association analysis. However, given the strong external validation in a wide range of data-sets and platforms, also including patients undergoing TKI treatment, we believe that our results are generalizable. Additional validity may be observed by its excellent performance when applied across platforms on RNA-Sequencing data, which would be of interest in future translation.

In summary, we report that a continuous quantitative scoring in ccRCC derived through conventional pathologic parameters yielded improved biological and clinical predictions over dichotomous subtype classifications, suggesting expression heterogeneity and corresponding biology is more suitably determined on a continuous scale. When applied to intratumoral regions, the CLEAR score suggests intertumoral heterogeneity generally exceeds intratumoral heterogeneity.

## Materials and Methods

### Public data collection

To evaluate ccRCC subtypes, public datasets were collected from GEO, EMBL-EBI and TCGA (The Cancer Geneome Atlas database) with the key words renal cell carcinoma or ccRCC. A total of 12 ccRCC microarray and RNA-seq data were collected (Supplementary Table S1). Consensus clustering sensitivity analysis was used to evaluate ccRCC subtypes from these 12 ccRCC datasets. Principal component analysis (PCA) and clustering typing with the well selection of reported ccRCC markers [13] were further applied to examine the appearance of ccRCC subtypes.

### Sample processing and data analysis

Expression profiles of 265 ccRCC samples were obtained from the Van Andel Research Institute. Baseline characteristics of patients are described in Supplementary Table S6. Gene expression profiles of these 265 samples were obtained using the HG-U133_Plus_2 platform. The resultant expression data was then summarized and normalized using the Robust Multi-array Average (RMA) method of the R “simpleaffy” Package (http://www.r-project.org). The expression data of 265 ccRCC samples were deposited in the Gene Expression Omnibus (GEO) under the accession number of GSE73731 (http://www.ncbi.nlm.nih.gov/geo/query/acc.cgi?token=udstaaakhbubjkt&acc=GSE73731)

### CLEAR score algorithm design, validation and screening of gene signatures

The analytic pipelines of CLEAR score algorithm design, validation and derivation of gene signatures are described and illustrated in Supplementary Methods and Supplementary Figure S2, S3. Comparison of CLEAR score and subtype model was also provided in supplemental Methods.

### Statistical analysis

We used the Fisher's exact test for investigating the enrichment and depletion of advanced levels of tumor grade, stage and size in different subgroups along the CLEAR scale. Spearman and Kendall’s tau correlation tests were used for evaluation of candidate signatures that contributed to the tumor prognosis ranking. Spearman correlation coefficient by exact test was used to analyze the correlation of three ranking queue. Kaplan-Meier survival test was used to assess the association of cancer-specific survival with the sample subgroups. Significance was determined using the log rank test. Other clinical covariates including age, tumor stage and tumor grade were compared to the outcome using univariate and multivariate Cox proportional hazards modeling. Log rank and likelihood ratio tests were done for multivariate modeling to assess statistical significance. All tests were performed using the R statistical computing package (http://www.r-project.org).

### Funding/Support and role of the sponsor

This study was supported by Institute of Bioengineering and Nanotechnology (Biomedical Research Council, Agency for Science, Technology and Research, Singapore).

## Acknowledgments

Tissue samples were provided by the Cooperative Human Tissue Network which is funded by the National Cancer Institute. We would like to thank the Van Andel Research Institute for support. We would like to sincerely thank Dr. Birgitta Sundelin, Dr. David Grignon and Dr. Shiro Baba for contributing tissue samples and Spectrum Health System Department of Pathology and Michigan Pathology Specialists for preparing and providing tissue samples. We would also like to thank Elizabeth Block, Jonathon Ditlev, and Michael Westfall for their assistance in running the samples.

